# Scarcity of scale-free topology is universal across biochemical networks

**DOI:** 10.1101/2020.09.16.299529

**Authors:** Harrison B. Smith, Hyunju Kim, Sara I. Walker

## Abstract

Biochemical reactions underlie all living processes. Like many biological and technological systems, their complex web of interactions is difficult to fully capture and quantify with simple mathematical objects. Nonetheless, a huge volume of research has suggested many real-world biological and technological systems – including biochemical systems – can be described rather simply as ‘scale-free’ networks, characterized by a power-law degree distribution. More recently, rigorous statistical analyses across a variety of systems have upended this view, suggesting truly scale-free networks may be rare. We provide a first application of these newer methods across two distinct levels of biological organization: analyzing a large ensemble of biochemical networks generated from the reactions encoded in 785 ecosystem-level metagenomes and 1082 individual-level genomes (representing all three domains of life). Our results confirm only a few percent of individual and ecosystem-level biochemical networks meet the criteria necessary to be anything more than super-weakly scale-free. Leveraging the simultaneous analysis of the multiple coarse-grained projections of biochemistry, we perform distinguishability tests across properties of individual and ecosystem-level biochemical networks to determine whether or not they share common structure, indicative of common generative mechanisms across levels. Our results indicate there is no sharp transition in the organization of biochemistry across distinct levels of the biological hierarchy - a result that holds across different network projections. This suggests the existence of common organizing principles operating across different levels of organization in biochemical networks, independent of the project chosen.

**Author Summary:** Fully characterizing living systems requires rigorous analysis of the complex webs of interactions governing living processes. Here we apply statistical approaches to analyze a large data set of biochemical networks across two levels of organization: individuals and ecosystems. We find that independent of level of organization, the standard ‘scale-free’ model is not a good description of the data. Interestingly, there is no sharp transition in the shape of degree distributions for biochemical networks when comparing those of individuals to ecosystems. This suggests the existence of common organizing principles operating across different levels of biochemical organization that are revealed across different network projections.

## Introduction

Statistical mechanics was developed in the 19th century for studying and predicting the behavior of systems with many components. It has been hugely successful in its application to those physical systems well-approximated by idealized models of non-interacting particles. However, real-world systems are often much more complex, leading to a realization over the last several decades that new statistical approaches are necessary to describe biological and technological systems. Among the most natural mathematical frameworks for developing the necessary formalism is network theory, which projects the complex set of interactions composing real systems onto an abstract graph representation [1–7]. Such representations are powerful in their capacity to quantitatively describe the relationship between components of complex systems and because they permit inferring function and dynamics from structure [8–12].

Network theory has been especially useful for studying metabolism. Metabolism consists of catalyzed reactions that transform matter along specific pathways, creating a complex web of interactions among the set of molecular species that collectively compose living things [13–17]. It is the collective behavior of this system of reactions that must be understood in order to fully characterize living chemical processes–counting only individual components (molecules) is inadequate. The structure of how those components interact with one another (via reactions) really matters: in fact it is precisely what separates organized biological systems from messy chemical ones [18–20].

Within the formalism of network theory, one of the simplest ways to capture insights into the global structure of a network is to analyze the shape of its degree distribution. A huge volume of research into various complex biological, technological and social networks has therefore focused on identifying the scaling behavior of the corresponding degree distributions for network projections describing those systems. One of the most significant results emerging from these analyses is that many networks describing real-world systems exhibit ostensibly “scale-free” topology [21–25], characterized by a power-law degree distribution. The allure of scale-free networks is in part driven by the simplicity of their underlying generative mechanisms, for example a power-law degree distribution can be produced by relatively simple preferential attachment algorithms [21], or to a lesser extent through optimization principles [26]. For truly scale-free networks the probability to find a node with degree *x* should scale as:

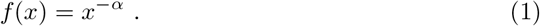

For numerous biological and technological systems, including metabolic networks, the scaling exponent, *α*, is reported with values in the range 2 *< α <* 3. The apparent ubiquity of scale-free networks across biological, technological and social networks has fueled some to conjecture scale-free topology as a unifying framework for understanding all such systems, with the enticing possibility these seemingly diverse examples could in reality arise from relatively simple, universal generating mechanisms [21, 25–28].

However, this story is far from complete. Recently developed statistical tests to rigorously examine whether observed distributions share characteristics with a power-law, or are instead more similar to other heavy tailed distributions, have revealed that true scale-free networks may not be as ubiquitous as previously supposed [29, 30]. These tests reveal that while it is superficially possible for a network to appear scale-free, more rigorous analysis can reveal a structure more similar to other heavy-tailed distributions such as the log-normal distribution, or even non heavy-tailed distributions like the exponential distribution [28–31].

The problem of characterizing the global structure of real-world systems is further compounded by the fact there are often many ways to coarse-grain a real system to generate a network representation, each corresponding to a different way for set of interactions to be projected onto a graph. For example, metabolic networks may be represented as unipartite or bipartite graphs, depending on whether one chooses to focus solely on the statistics over molecules (or reactions) and their interactions (requiring a unipartite representation) or instead to include both molecules and reactions as explicit nodes in the graph (where molecules and reactions represent two classes of nodes in a bipartite representation) [32–34]. These graphs can have different large-scale topological properties, even when projected from the same underlying system. This raises the question of determining which projection to analyze, and whether or not a real-world system should be considered “scale-free” if only some of its network projections exhibit power-law degree distributions. Broido and Clauset recently developed a methodology to compare the degree distributions of network projections of different complexities, classifying the degree to which they are scale-free on a scale from “Not scale-free” all the way to “strongest” [30]. This provides a framework for statistically analyzing many projections of a given system to determine how well scale-free structure describes the real underlying system when projected onto its different coarse-grained representations.

Herein, we build from the work of Broido and Clauset with specific application to the problem of characterizing biochemical systems. A novelty in our approach is recognizing that in order to really understand the structure of real-world biological (and technological) systems, the relevant scale(s) for performing such analysis must also be considered. In particular, many biological and technological systems are hierarchical, with networks describing interactions across multiple levels. For example, one may study the biochemistry of individual species, but ultimately the function of an individual in a natural system depends on a complex interplay of interactions among the many species comprising its host ecosystem. In this way, biochemistry is hierarchically organized into individuals and ecosystems. Indeed, much discussion about universal properties of life has shifted focus from individuals to ecosystems as the relevant scale best capturing the regularities of biological organization [35, 36]. It is unclear at present whether analysis of biochemical networks at the level of individuals or ecosystems will best uncover their structure and permit identifying generative mechanisms for biology, or whether all levels must be considered simultaneously.

In what follows, we perform statistical analysis of an ensemble of biochemical systems generated from 785 ecosystem-level metagenomes and 1082 individual-level genomes (representing all three domains of life). Our results include the first analysis of scale-free network structure for the different projections of ecosystem-level biochemistry, significantly expanding on on earlier work focusing on the large-scale organization of individual metabolic networks only [13, 29, 30, 32–34]. Like Broido and Clauset, we consider all possible projections of biochemical systems to graphs simultaneously, whereas most prior work on the organization of biochemistry has only considered one or at most a few projections [17, 37–40]. We find a majority of biochemical networks are not scale-free, independent of projection or level of organization. We also demonstrate how the network properties analyzed herein can be used to distinguish individual and ecosystem level networks, and find that independent of projection, individuals and ecosystems share very similar structure. These results have potentially deep implications for identifying underlying rules of biochemical organization at both the individual and ecosystem-level by providing constraints on whether the same or different generative mechanisms could operate to organize biochemistry across multiple scales.

## Results

Utilizing the framework developed by Broido and Clauset [30], we used the full set of biochemical reactions encoded in each genome and metagenome to construct eight distinct network representations of each respective biochemical system. This resulted in 8656 network projections for the 1082 individual-level biochemical datasets, and 6280 network projections for the 785 ecosystem-level biochemical datasets. Each representation corresponds to a different coarse-graining of the underlying system of reactions (i.e. the underlying dataset) (**Fig. 1**). We determine whether or not these datasets are scale-free, and analyze the aspects of them, and their diverse projections, that tend to lend themselves to be more or less scale-free. The alternative distributions that we compare to the power-law are: The exponential distribution, the log-normal distribution, the stretched exponential distribution, and the power-law distribution with a cutoff (see [29, 30] for more details on these distributions).

**Figure 1.**
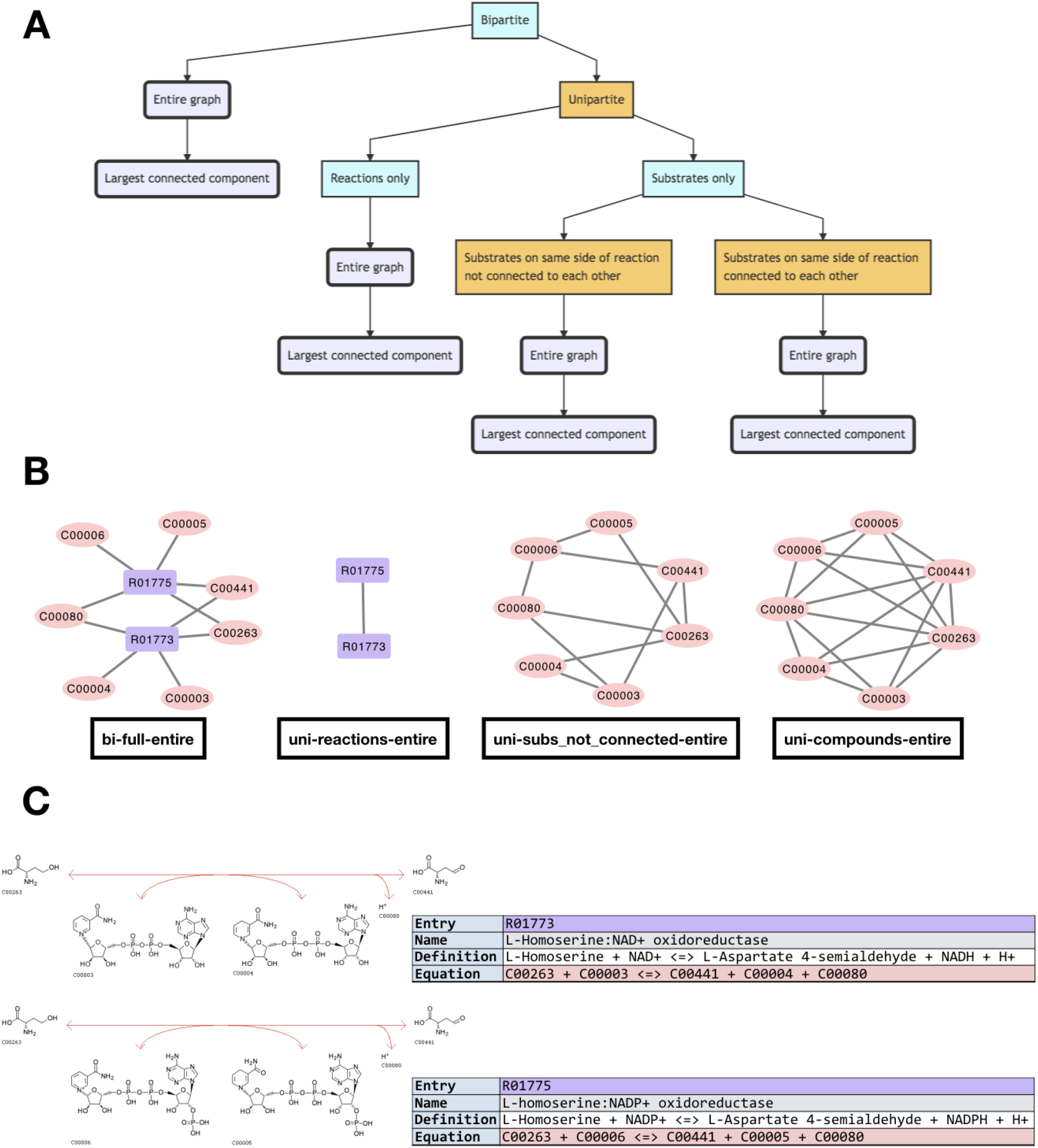
How biochemical datasets are decomposed into network projections. **(A)** Networks are generated from the set of reactions encoded in each genome/metagenome starting from a bipartite representation, and projecting different combinations of attributes. The bold, rounded flowchart nodes show the result of each combination of projections applied in this study. **(B)** The different network projection types of a simple example dataset, composed of two KEGG reactions: R01773 & R01775. The nomenclature used in this paper’s figures is below each network visualization (in this example the entire graph is the same as the largest connected component). **(C)** How the reactions used in the network visualization example above appear in the KEGG database [41–43]

We first classified each dataset in terms of how scale-free it is. A dataset is classified as: *Super-Weak* if for at least 50% of network projections, none of the alternative distributions are favored over the power-law; *Weakest* if for at least 50% of network projections, the power-law hypothesis cannot be rejected (*p ≥* 0.1); *Weak* if it meets the requirements of the Weakest set, and there are at least 50 nodes in the distribution’s tail (*n*_tail_ *>* 50); *Strong* if it meets the requirements of both the Super-Weak and Weak set, and that the median scaling exponent is between two and three 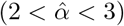; and *Strongest* if it meets the requirements of the Strong set for at least 90% of graphs, rather than 50%, and for at least 95% of graphs none of the alternative distributions are favored over the power-law.

Our results indicate most biochemistry at the individual and ecosystem-level is characterized by networks that are “super-weakly” scale-free (**Fig. 2**). That is, while the power-law is better than other models for fitting the shape of their degree distributions, the power-law is not itself a good model. When doing a goodness-of-fit test, we find that the majority of network representations of each genomic/metagenomic dataset have *p <* 0.1, indicating there is less than a 10% chance that our data is truly power-law distributed. This effectively rules out the possibility that our data is drawn from a power-law shaped degree distribution, despite the fact that, when compared to other distributions through log-likelihood ratios, 99% of all datasets do not favor alternative heavy tail distributions for the majority of their network-projections (**Fig. 3**, top row).

**Figure 2.**
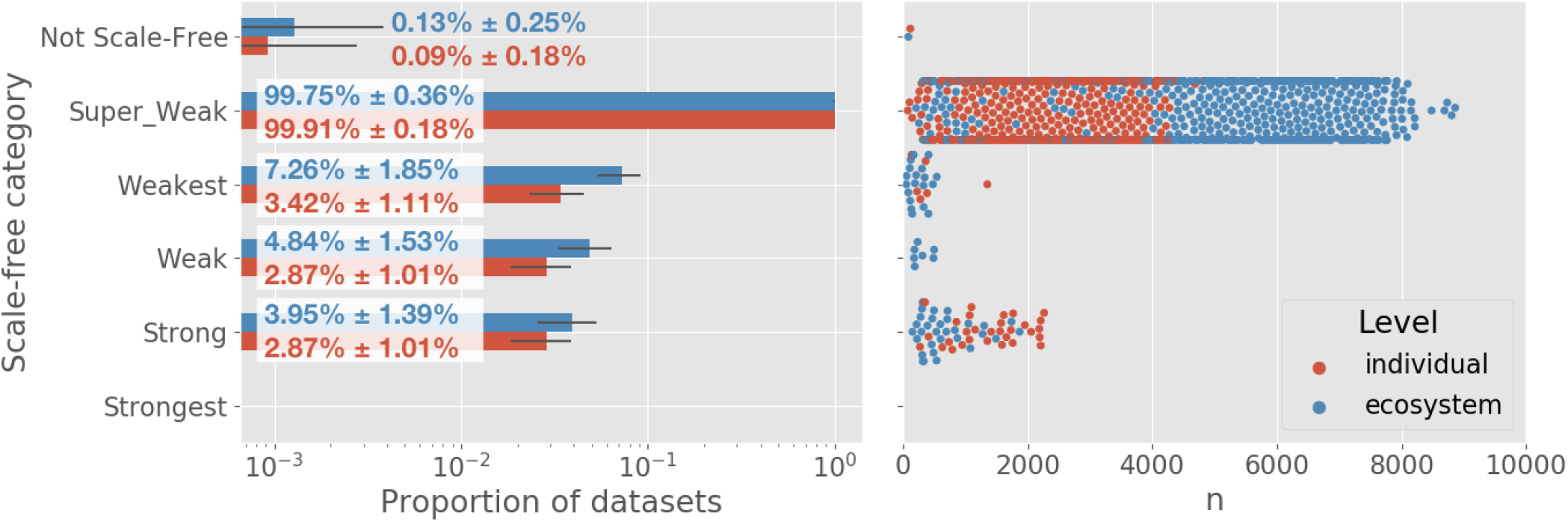
The vast majority of individual and ecosystem level networks are not “scale-free”. Left: Most datasets are super weak, indicating that when compared to other models, a power-law distribution is a better fit. However, the power-law distribution is not a “good” fit for most dataset network representations. No networks meet the “Strongest” criteria defined by Broido and Clauset al. [30]. Overlaid values show the percent of networks of each level which fall into each category, *±*2SD. Right: The relationship between scale-freeness and largest network size across projections (*n*). All datasets containing networks larger than approximately 2100 nodes have degree distributions that rule out fitting well to a power-law.

**Figure 3.**
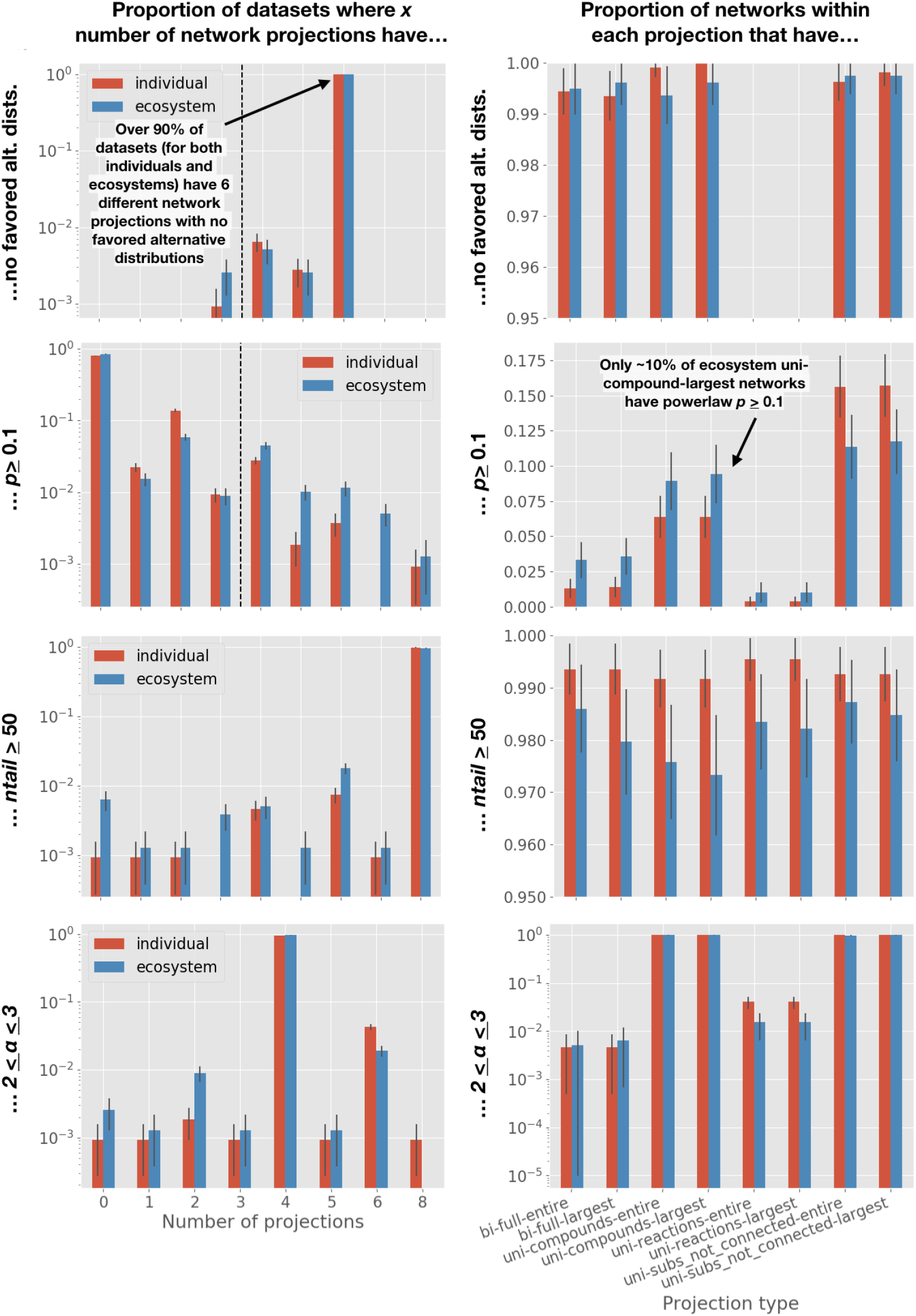
The number of network projections within each dataset which meet some scale-free criteria. Left column: The number of network projections within each dataset which meet some scale free-criteria, where each dataset falls into one of nine bins. Normalized to total number of datasets in a level. Criteria from top to bottom: No alternative distributions favored over power-law in log-likelihood ratio (1st row); *p ≥* 0.1 (2nd row); *n*_*tail*_ *>* 50 (3rd row); 2 *< α <* 3 (4th row). Dashed lines show: the cutoff for number of networks in a dataset required to meet the threshold criteria for “Super-Weak” (1st row), and “Weakest” (2nd row). Right column: The number of network projections, across all datasets, which meet some scale-free criteria, binned by projection type. Normalized to the total number of each projection within a level. Criteria same as left column. Red bars indicate individual-level datasets/networks, and blue bars indicate ecosystem-level datasets/networks. Black error bars show *±*2SD.

### Where biochemical systems succeed and fail scale-free classifications

#### Goodness-of-fit *p*-value

The “weakest” requirement for a scale-free network introduced by Broido and Clauset stipulates over 50% of a dataset’s network-projections must have a power-law goodness-of-fit *p ≥* 0.1. For both individuals and ecosystems, only 6% of network-projections meet this requirement (**Fig. 4**, left column). This goodness-of-fit *p*-value requirement is the most restrictive of all scale-free requirements.

**Figure 4.**
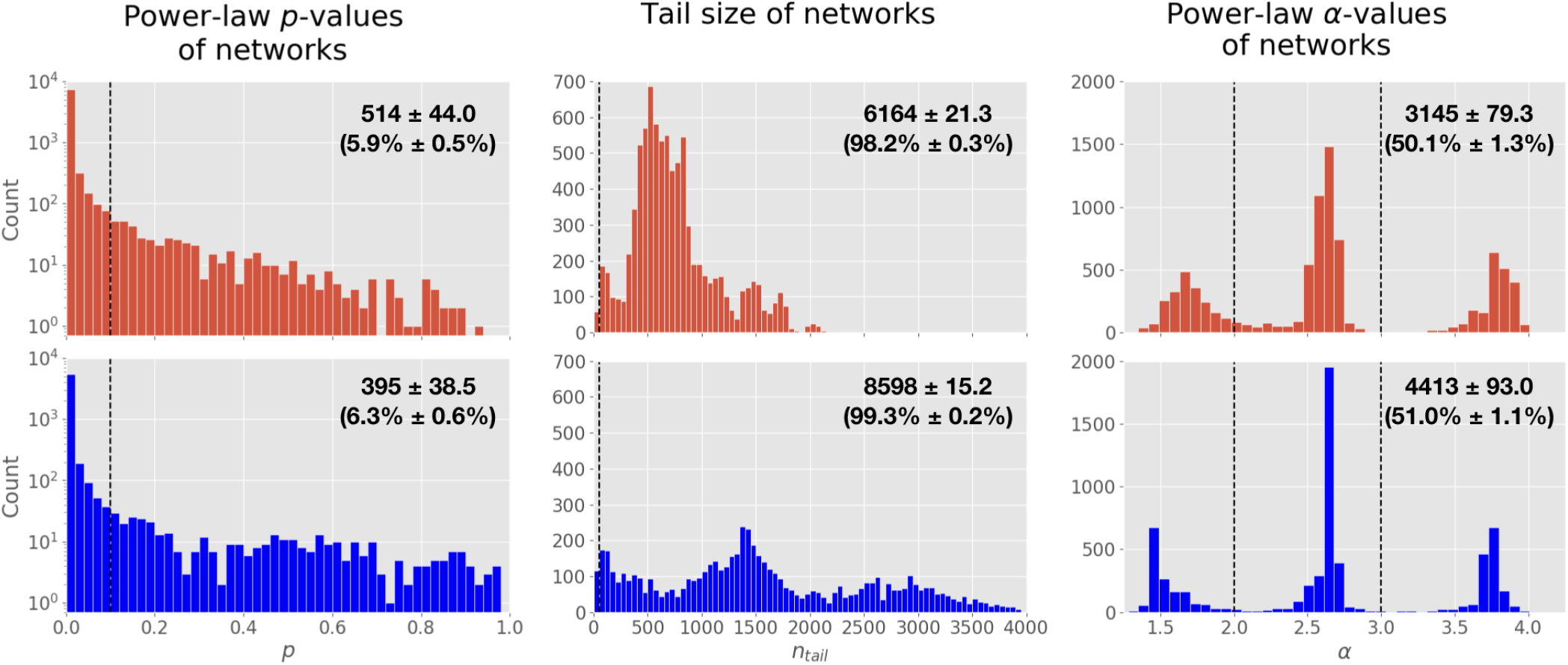
The distribution of *p*-values, tail-sizes, and power-law alpha values for biochemical network degree distributions, over all network projections. Left column: The goodness-of-fit *p*-values of networks. When *p ≥* 0.1 (dashed line), it indicates that there is at least a 10% chance of the power-law distribution being a plausibly good fit to a network’s degree distribution. Center column: Tail size of networks. When *n*_*tail*_ ≥ 50 (dashed line), it indicates that the tail of distribution is large enough to reliably fit. Right column: Power-law exponent *α* values of networks. When 2 *< α <* 3 (between dashed lines), it indicates that a network meets the criteria of having a power-law exponent which falls into scale-free territory. The top row shows distributions for individuals in red. The bottom row shows distributions for ecosystems in blue. Insets indicate the number (and percent) of networks which meet the criteria, *±*2SD.

#### Tail size

Setting aside the fact each subsequent scale-free requirement builds on the requirement(s) of the preceding one, we find 98% of individual networks and 99% of ecosystem networks *do* meet the requirement of *n*_*tail*_ *>* 50 for a scale-free degree distribution (**Fig. 4**, center column).

#### The power-law exponent, *α*

Only 50% of individual-level networks and 51% of ecosystem-level networks meet the requirement that 2 < *α* < 3 for their degree distribution. The goodness-of-fit *p* value requirement, followed by the requirement constraining values of *α*, are the most restrictive when determining whether a biochemical network’s degree distribution should be considered scale free (**Fig. 4**, right column).

### Meeting the threshold for scale-free classification is dependent on the network representation

We find the results of each requirement listed above for classifying topology as scale-free differ across the eight network projection types for each dataset. Unsurprisingly, for most requirements, there exists a minute difference between the values observed for the largest connected component and entire graph of a given network projection type (**Fig. 3**, right column). Depending on the measure, there is a noticeably larger difference between the major network projection types, e.g., between bipartite, unipartite-reactions, unipartite-compounds (where all substrates participating in the same reaction are connected), and unipartite-compounds (where substrates on the same side of a reaction are not connected) (**Fig. 3**, right column).

#### Comparing to alternative distributions

Over 99% of individual and ecosystem-level datasets have 6 projections which do not favor any other distribution over the power-law (**Fig. 3**, top row, left column). No datasets have more than 6. The other two projections nearly always favor at least one other distribution over the power-law distribution–either the log-normal, exponential, stretched exponential, or power-law with exponential cutoff (**Fig. 3**, top row, right column). There are only 3 of the 6280 ecosystem-level network projections (across the 785 ecosystem-level datasets) that do not favor at least one of the alternative distributions. Oftentimes all four are favored over the power-law distribution (**Fig. S6**, rows 3-4). These results are identical, within 95% confidence, for both individuals and ecosystems.

#### Goodness-of-fit *p*-value

Out of all datasets, 80% of individuals and 84% of ecosystems have only a single projection type with *p* ≥ 0.1 for a power-law fit to their degree distribution. This indicates the majority of datasets would still not meet the “weakest” requirement for scale-free even with a threshold that lowered the percent of a dataset’s projections needed to 25% (2 networks) instead of 50% (4 networks) (**Fig. 3**, 2nd row, left column). The unipartite projection where substrates on the same side of a reaction are not connected (unipartite-subs not connected) was the most likely to satisfy *p* ≥ 0.1. For the two unipartite-compound projections, the difference between individuals and ecosystems is within the error. The unipartite-reaction projections were the least likely to satisfy *p* ≥ 0.1, which is consistent with the observation that these networks always favor an alternative distribution as a better fit to the data than the power-law (**Fig. 3**, 2nd row, right column). As we initially reported, the majority of datasets do not meet the *p*-value threshold for being considered scale-free, although ecosystems-level datasets are more likely to meet the threshold.

#### Tail size

Out of all datasets, 98% of individuals and 96% of ecosystems meet *n*_*tail*_ *>* 50 for all projection types (**Fig. 3**, 3rd row, left column). For 7 of the projection types, there is no difference between individuals and ecosystems, within 95% confidence (**Fig. 3**, 3rd row, right column).

#### The power-law exponent, *α*

Out of all datasets, 95% of individuals and 97% of ecosystems meet 2 *< α <* 3 for 4 of 8 projection types (**Fig. 3**, bottom row, left column). The two types of unipartite-compound networks contribute to the datasets which meet the alpha-range requirement the majority of the time. That is, chances are if a dataset has at least 4 projection types meeting 2 *< α <* 3, two of them are going to be unipartite-compound network projections (**Fig. 3**, bottom row, right column). The results are similar for both individuals and ecosystems.

#### Correlation of results between projections

Because 8 different network projections are derived from a single biochemical dataset, there is reason to expect the proportions of each projection type meeting any given scale-free criteria are correlated. We therefore constructed a Pearson correlation matrix to test whether there are correlations between projections (**Fig. S7**). Unsurprisingly, we find that values from projections of a network’s LCC and entire graph are highly correlated. All types of unipartite compound networks tend to be correlated. Values across many other projection types are barely correlated for the *p*-value and *n*_*tail*_ criteria. Ecosystems tend to show more correlation, across all projection types, than individuals.

### Distinguishing individuals and ecosystems based on their degree distributions

#### Multinomial regression

We used multinomial regression on network and degree distribution data from the above analyses to attempt to distinguish individuals from ecosystems. Most measures cannot reliably distinguish between these two levels of organization, with only network size and network tail size data distinguishing the two levels better than chance. Using only network size, ecosystems could be correctly identified in test data 72.23% of the time, whereas individuals could be correctly identified 85.33% of the time (**Fig. 5**, left columns). When normalizing other measures to network size, the only one that improved in distinguishing individuals and ecosystems to be better than chance was *dexp* (**Fig. S8**). This is a measure of which type of distribution is favored (or neither) when doing a log-likelihood ratio test between the power-law and exponential distribution.

**Figure 5.**
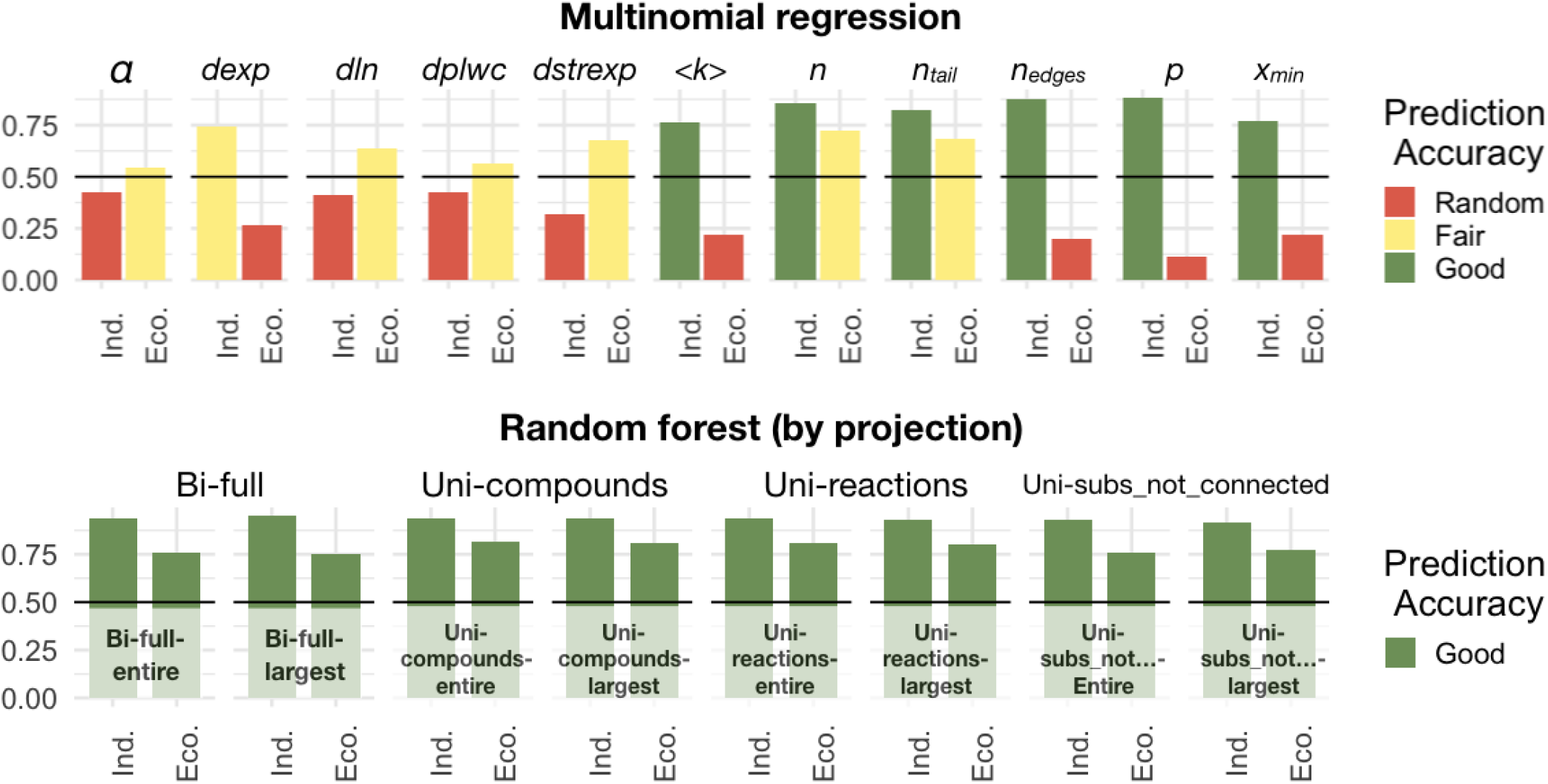
Predicting individuals and ecosystems from degree distribution data using multinomial regression vs. random forest. Each subplot shows the accuracy of using a particular network or statistical measure to predict whether that network data came from an biological individual or ecosystem. The top plots show prediction accuracy from using multinomial regression across all network projection types, and the bottom plots show prediction accuracy using random forest on each type of projection. The random forest classifier is much better at predicting individuals and ecosystems correctly from network data, even without direct access to network size. All random forest predictions have an accuracy of at least 75% across all projection types. Subplots measures are: power-law alpha value; log-likelihood result from power-law vs. exponential; log-likelihood result from power-law vs. log-normal; log-likelihood result from power-law vs. power-law with exponential cutoff; log-likelihood result from power-law vs. stretched exponential; the network mean degree; network node size; degree distribution tail size; network edge size; the *p*-value of the goodness-of-fit test for the power-law model; cutoff degree value for network tail. These are the only predictors used in the random forest classifier. Prediction accuracy is random if *≤* 50%, Fair if *>* 50%, and Good if *≥* 75%.

**Figure 6.**
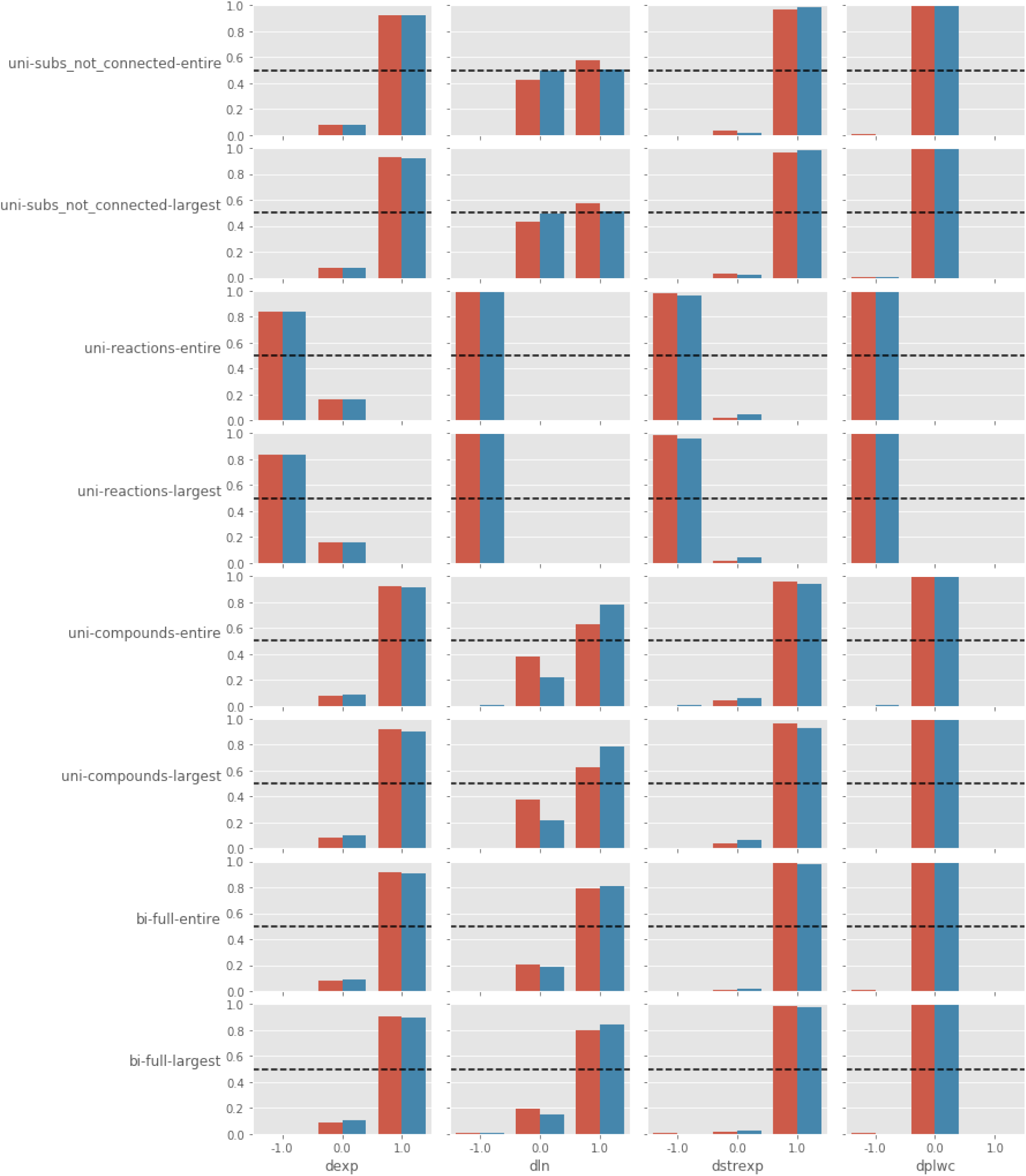
How alternative distributions compare to the powerlaw across each network projection type. The proportion of network projections, across all datasets, that favor either the power-law distribution (1.0), an alternative distribution (−1.0), or are inconclusive (0.0). Each row shows a different network projection type. Each column is a different distribution with which the power-law is being compared to. From left to right is the exponential; log-normal; stretched exponential; and power-law with cutoff. Dashed line is constant at proportion = 0.5 across all subplots. Red bars indicate individual-level networks, and blue bars indicate ecosystem-level networks.

**Figure 7.**
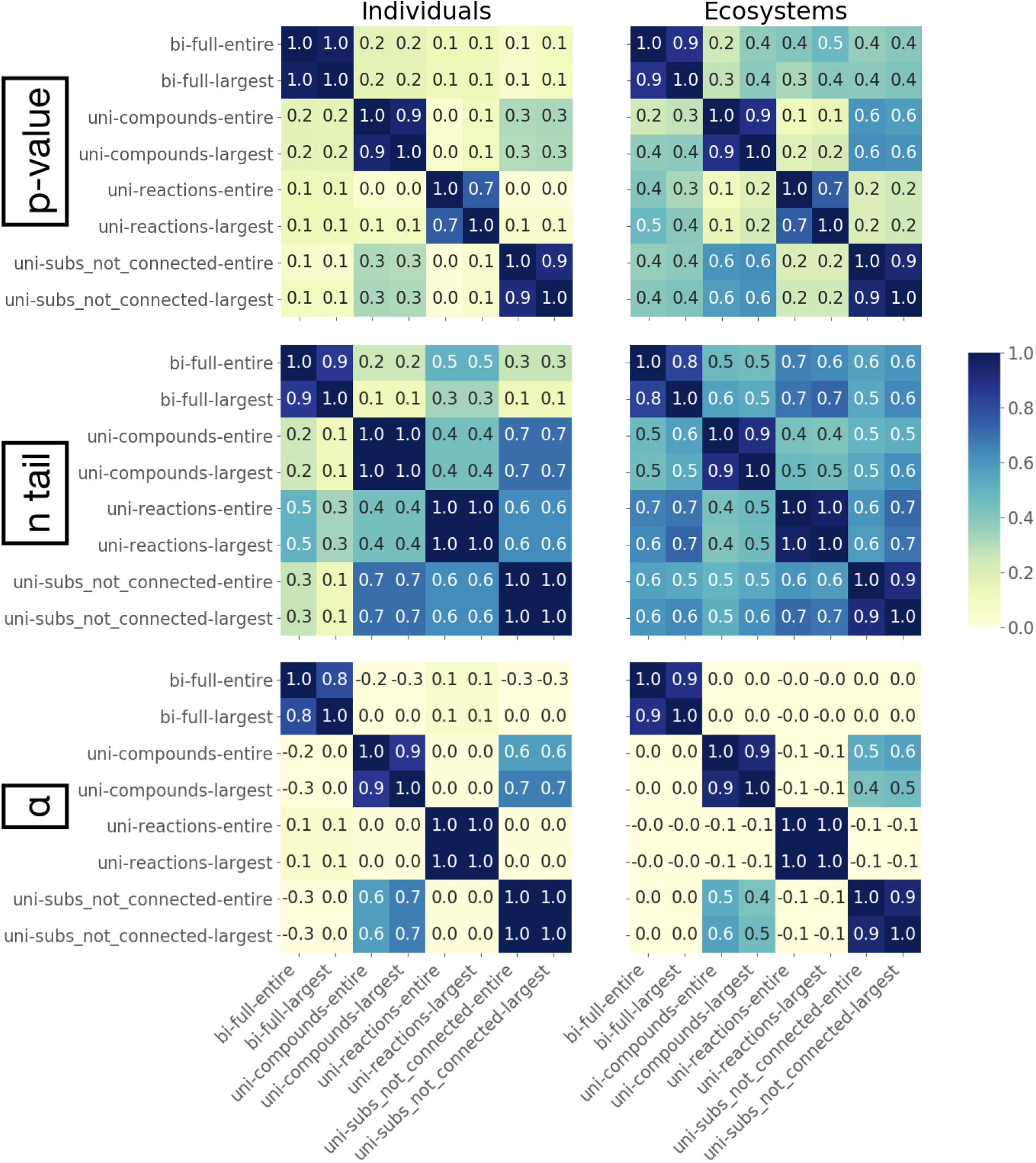
Correlations between network projections which meet scale-free criteria. Correlation matrix heatmaps show type how different types of network projections correlate in their proportions of networks which meet some scale-free criteria. Rows are for each of the different scale-free criteria (*p*-value, *n*_*tail*_ and *α*), and columns are for individual and ecosystem-level networks. Heatmaps show the correlation between values for each projection type, where the values are of the proportion of networks which meet the scale-free threshold criteria of: *p* ≥ 0.1 (top row); *n*_*tail*_ *>* 50 (center row); 2 *< α <* 3 (bottom row). Values from projections of a network’s LCC and entire graph are highly correlated. All types of unipartite compound networks tend to be correlated. Values across many other projection types are barely correlated for the *p*-value and *n*_*tail*_ criteria. Ecosystems tend to show more correlation, across all projection types, than individuals. log-likelihood result from power-law vs. stretched exponential; the log-likelihood of the power-law model; the network mean degree; network node size; degree distribution tail size; network edge size; cutoff degree value for network tail. Prediction accuracy is random if *≤* 50%, Fair if *>* 50%, and Good if *>* 75%.

**Figure 8.**
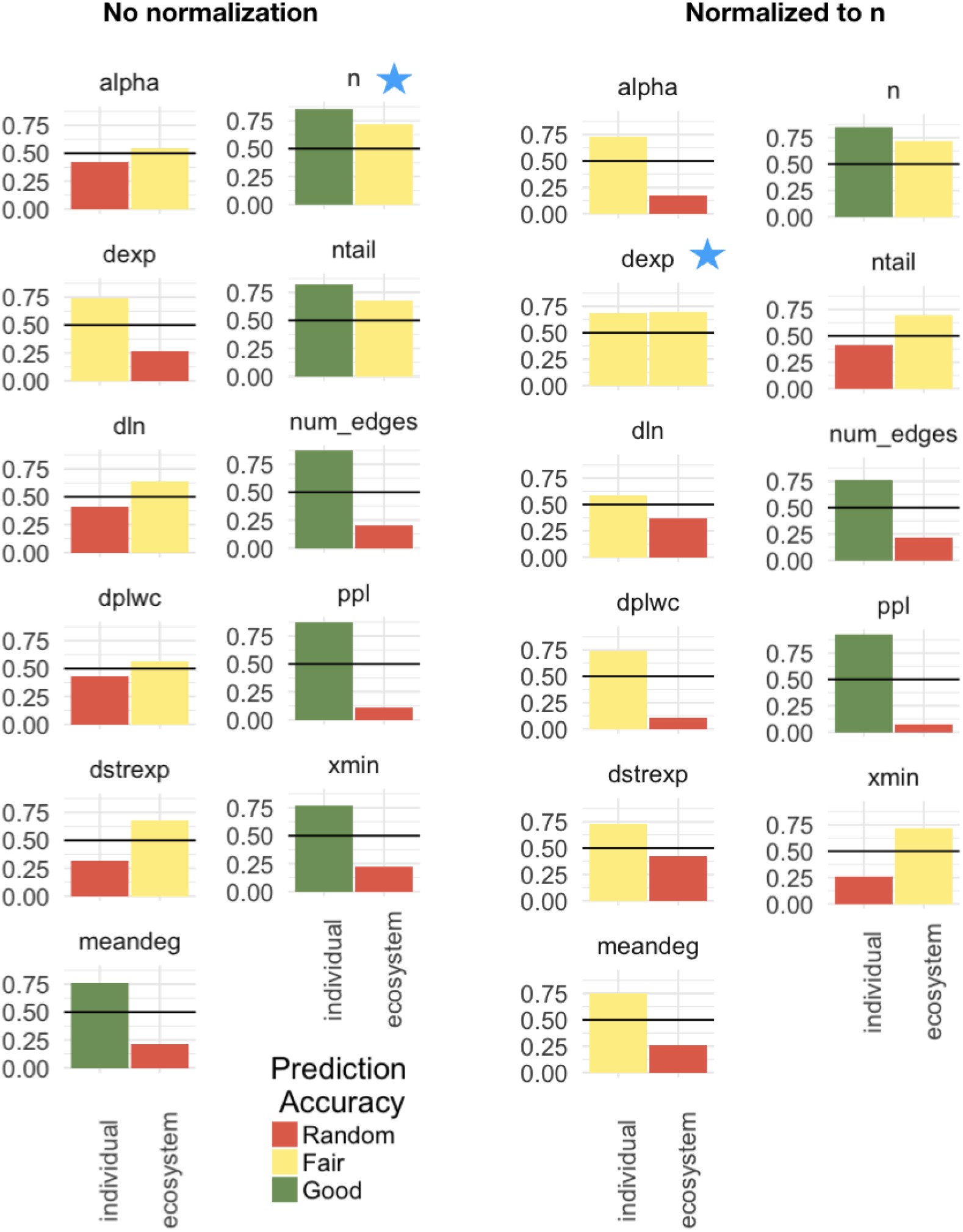
Predicting individuals and ecosystems from degree distribution data using multinomial regression. Each subplot shows the accuracy of using a particular network or statistical measure to predict whether that network data came from an biological individual or ecosystem. The subplots in the right column are the accuracy of using a measure after being normalized to network size. Unsurprisingly, network size is by far the best way to accurately predict whether data comes from an individual or ecosystem (left blue star). Once normalized to size, whether or not a degree distribution favors an exponential fit compared to a power-law fit becomes a decent predictor (right blue star). Subplots measures are: power-law alpha value; log-likelihood result from power-law vs. exponential; log-likelihood result from power-law vs. log-normal; log-likelihood result from power-law vs. power-law with exponential cutoff; log-likelihood result from power-law vs. stretched exponential; the network mean degree; network node size; degree distribution tail size; network edge size; the *p*-value of the goodness-of-fit test for the power-law model; cutoff degree value for network tail. Prediction accuracy is random if *≤* 50%, Fair if *>* 50%, and Good if *>* 75%.

**Table 1.**
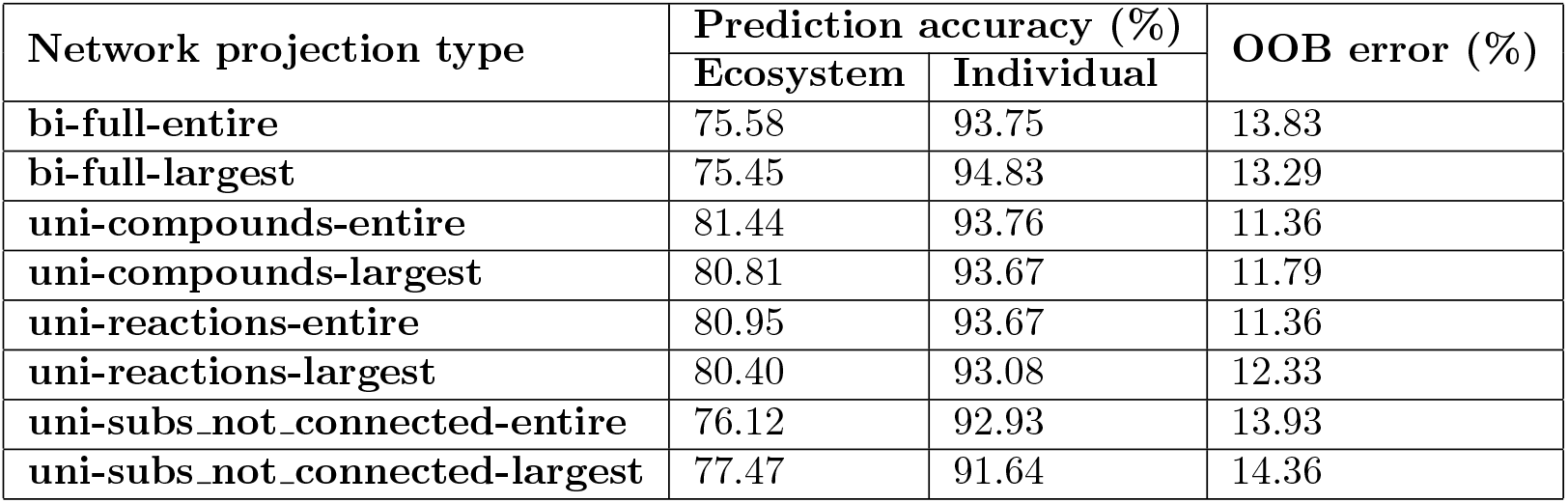
Random forest accuracy by network projection type.

#### Random Forest

Random forest classifiers are a supervised machine learning technique that use decision trees to make classifications. When using random forests to try and distinguish individuals and ecosystems based on network and distribution data, we find ecosystems can be correctly predicted 87.01% of the time, and individuals can be correctly predicted 95.82% of the time (Out of bag, OOB, error rate is 7.91%). However, the size of the network and size of the degree distribution tail once again are the best relative predictors. Without network size and tail size, the prediction accuracy drops to 79.27% for ecosystems and 94.81% for individuals (OOB error rate of 11.80%). When doing random forest classification by projection type, the prediction accuracies are still above 75% for ecosystems and 91% for individuals across all projections, which is better than multinomial regression models even when information about network size is included (**Fig. 5**, left columns; **Table S1**). Mean degree was the best predictor across all network projection types.

## Discussion

Our results indicate biochemical systems across individuals and ecosystems are, at best, only weakly scale-free. This is revealed by studying all possible projections of biochemical systems in tandem: only six of the eight network projection types analyzed favor power-law distributions over alternatives and in all cases the power-law is not itself a good fit to the data. Nonetheless, we can conclude individuals and ecosystems both share qualitatively similar degree distribution characteristics, and while this is a very coarse-grained measure of network structure, it suggests the possibility of shared principles operating across levels of organization to architect biochemical systems. The random forest distinguishability analyses demonstrate how using a combination of all the results of scale-free analyses completed in this paper can predict, better than chance, whether the data comes from individuals or ecosystems. Individuals are perhaps more tightly constrained in their network structure, based on being able to more accurately predict them based on simple network characteristics. Whether or not this structure is truly a universal property of life’s chemical systems is more difficult to conclude. Based on the sample sizes, we are confident our results hold over the population of genomes and metagenomes in the JGI and PATRIC databases. However, the observed scaling is only reflective of biology universally if the databases are unbiased in sampling from all of biology on Earth, and this is impossible to know with certainty (see *e*.*g*. proposals of ‘shadow life’ and reports of missing biota [44, 45]). Nonetheless, the fact that multiple levels and multiple projections of biochemistry reveal common structure suggests universal principles may be within reach if cast within an ensemble theory of biochemical network organization (see *e*.*g*. also [36]).

Achieving such a theory requires recognizing that, unlike simple physical systems where statistics over individual components is sufficient to describe and predict their behavior, biological and technological systems require additional information about the structure of interactions among their many components. This is well-known, but how to project this structure onto simple mathematical objects that can be quantifiably characterized and compared remains a central problem of complex systems science. In physics, the relevant coarse-graining procedure is well understood, but we are not so far in understanding living processes: the first hurdle we must traverse is to identify the proper coarse-grained network representations for analysis. Existing literature cautions against using unipartite network projections, as it is argued they can lead to “wrong” interpretations of system properties such as degree in biochemical networks [34, 46]. We find instead that whether or not this conclusion should be drawn is highly dependent on the particular characteristics of degree or the degree distribution under consideration. For example, all network projection types, aside from unipartite reaction networks, favor power-law degree distributions over other heavy-tailed alternatives (**Fig. 3**, top row). For power-law *α* ranges 2 *< α <* 3, bipartite networks show similar results to the unipartite reaction networks of individuals, but different results for ecosystems and unipartite compound networks (**Fig. 3**, fourth row). Almost all projections show differing results for meeting the scale-free *p*-value cutoff (**Fig. 3**, second row). While other literature [32, 33] has advocated for unipartite networks (with all compounds participating in a reaction connected–called *uni-compounds* here), we find that these networks overestimate power-law goodness-of-fit *p*-values and *α* values compared to reaction and bipartite networks (**Fig. 3**). The similarities and differences in the structure of different projections provides insight into the actual structure of the underlying system of interest. Given that there is no obvious answer for whether a system is scale-free, we advocate for studying all projections possible: regardless of whether or not a given projection is scale-free, all projections provide insights into the structure of the underlying system. In physics we have become accustomed to one unique coarse-grained descriptor providing insight into the structure of a system. It may be that to really understand complex interacting systems, such as the systems of reactions underlying all life on Earth, we must forget the allure of simple, singular models. Instead, to characterize the regularities associated with living processes, we should perform statistical analyses over many projections to determine what features are truly universal.

## Materials and Methods

### Obtaining biological data

Bacteria and Archaea data were obtained through PATRIC [47]. Starting with the 21,637 bacterial genomes available from the 2014 version of PATRIC, we created a parsed dataset by selecting one representative genome containing the largest number of annotated ECs from each genus. Unique genera (genera only represented by a single genome) were also included in our parsed data. Uncultured/candidate organisms without genera level nomenclature are left in the parsed dataset. This left us with 1152 parsed bacteria, from which we chose 361 randomly to use in this analysis. Starting with 845 archaeal genomes available from the 2014 version of PATRIC, we randomly chose 358 to use in this analysis. Enzyme Commission (EC) numbers associated with each genome were extracted from the `ec number` column of each genome’s `.pathway.tab` file.

Eukarya and Metagenome data were obtained through JGI IMG/m [48]. All 363 eukaryotic genomes available from JGI IMG/m as of Dec. 01, 2017 were used. Starting with the 5586 metagenomes available from JGI IMG/m as of June 20, 2017, 785 metagenomes were randomly chosen for this paper’s analyses. Enzyme Commission (EC) numbers associated with each genome/metagenome were extracted from the list of *Protein coding genes with enzymes*, and metagenome EC numbers were obtained from the *total* category. All JGI IMG/m data used in this study were sequenced at JGI.

Because each EC number corresponds to a unique set of reactions that an enzyme catalyzes, the list of EC numbers associated with each genome and metagenome can be used to identify the reactions that are catalyzed by enzymes coded for in each genome/metagenome. We use the Kyoto Encyclopedia of Genes and Genomes (KEGG) ENZYME database to match EC numbers to reactions, and the KEGG REACTION database to identify the substrates and products of each reaction [41–43]. This provides us with a list of all chemical reactions that a genome/metagenome’s enzymes can catalyze.

### Generating Networks

Each genomic/metagenomic dataset is used to construct eight representations of biochemical reaction networks. We refer to each type of representation as a “network projection type” throughout the text:

1. *Bipartite graph with reaction and compound nodes*. A compound node *C*_*i*_ is connected to a reaction node *R*_*i*_ if it is involved in the reaction as a reactant or a product. Abbreviated in figures as *bi-full*.
2. *Unipartite graph with compound nodes only*. Two compound nodes *C*_*i*_ and *C*_*j*_ are connected if they are both present in the same reaction. A reaction’s reactant compounds are connected to each other; a reaction’s product compounds are connected to each other; and a reaction’s reactant and product compounds are connected. Abbreviated in figures as *uni-compounds*.
3. *Unipartite graph with reaction nodes only*. Two reaction nodes *R*_*i*_ and *R*_*j*_ are connected if they involve a common compound. Abbreviated in figures as *uni-reactions*.
4. *Unipartite graph with compound nodes only (alternate)*. Two compound nodes *C*_*i*_ and *C*_*j*_ are connected only if they are both present on opposite sides of the same reaction. A reaction’s reactant compounds are *not* connected to each other; a reaction’s product compounds are *not* connected to each other; but a reaction’s reactant and product compounds are connected. Abbreviated in figures as *uni-subs not connected*.

There exists a version of each of these four network construction methods for the largest connected component (LCC), and for the entire graph, yielding a total of eight network projections for each dataset (**Fig. 1**). These network projection types are signified in the figured by appending *-largest* and *-entire* to the network projection abbreviations. Some datasets may yield identical networks for their LCC and entire graph, if there is exists only a single connected component.

### Assessing the power-law fit on degree distributions

As defined in Clauset, 2009 [29], a quantity *x* obeys a power law if it is drawn from a probability distribution

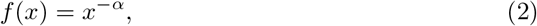

where *α*, the exponent/scaling parameter of the distribution, is a constant. In order to estimate *α*, we follow the methods described in Clauset, 2009 [29], and use an approximation of the discrete maximum likelihood estimator (MLE)

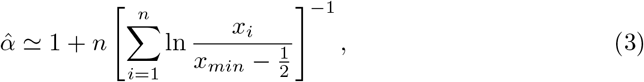

where *x*_*min*_ is the lower bound of power-law behavior in our data, and *x*_*i*_, *i*=1,2,…,n, are the observed values *x* such that *x*_*i*_ ≥ *x*_*min*_. The standard error of our calculated *α* is given by

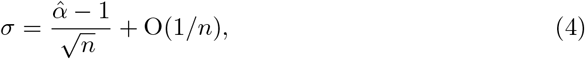

where the higher-order correction is positive [29]. Because many quantities only obey a power-law for values greater than some *x*_*min*_, the optimal *x*_*min*_ value must be calculated. The importance of choosing the correct value for *x*_*min*_ is discussed in detail in Clauset et al, 2009 [29]. If it is chosen too low, data points which deviate from a power-law distribution are incorporated. If it is chosen too high, the sample size decreases. Both can change the accuracy of the MLE, but it is better to err too high than too low.

In order to determine *x*_*min*_, we use the method first proposed by Clauset et al., 2007 [49], and elaborated on in Clauset et al., 2009 [29]: we choose the value of *x*_*min*_ that makes the probability distributions of the measured data and the best-fit power-law model as similar as possible above *x*_*min*_. The similarity between the distributions is quantified using the Kolmogorov-Smirnov or KS statistic, given by

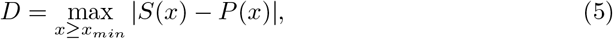

where *S*(*x*) is the cumulative density function (CDF) of the data for the observations with value at least *x*_*min*_, and *P* (*x*) is the CDF for the power-law model that best fits the data in the region *x* ≥ *x*_*min*_. Our estimate of *x*_*min*_ is the one that minimizes D.

We used the github respository made available in Broido and Clauset [30] to determine the optimal *x*_*min*_ of all our degree distributions, and to subsequently calculate the MLE in order to determine the *scaling exponent α* and the *standard error* on *α, σ* [50].

A power-law can always be fit to data, regardless of the true distribution from which it is drawn from, so we need to determine whether the power-law fit is a good match to the data. We do this by sampling many synthetic data sets from a true power-law distribution, recording their fluctuation from power-law form, and comparing this to similar measurements on the empirical data in question. If the empirical data has similar form to the synthetic data drawn from a true-power law distribution, then the power-law fit is plausible. We use the KS statistic to measure the distance between distributions.

We use a goodness-of-fit test to generate a *p*-value which indicates the plausibility of a hypothesis. The *p*-value is defined as the fraction of the synthetic distances that are larger than the empirical distance. If *p* is large (close to 1), then the difference between the empirical data and the model can be attributed to statistical fluctuations alone; if it is small, the model is not a plausible fit to the data [29]. We follow the methods in Clauset et al., 2009 [29]–and implement them with the github package used in Broido and Clauset [30]–to generate synthetic datasets and measure the distance between distributions. Following these methods, we chose to generate 1000 synthetic datasets in order to optimize the trade-off between having an accurate estimation of the *p*-value and computational efficiency. If *p* is small enough (*p <* 0.1) the power law is ruled out. Put another way, it is ruled out if there is a probability of 1 in 10 or less that we would by chance get data that agree as poorly with the model as the data we have [29]. However, measuring a *p* ≥ 0.1 does not guarantee that the power-law is the most likely distribution for the data. Other distributions may match equally well or better. Additionally, it is harder to rule out distributions when working with small sample sizes.

A better way to determine whether or not data is drawn from a power-law distribution is to compare its likelihood of being drawn from a power-law distribution directly to a competing distribution [29, 51]. We use the exponential, stretched-exponential, log-normal, and power-law-with-cutoff distributions as four competing distributions to the power-law. While we cannot compare how the data fits between every possible distribution, comparing the power-law distribution to these four similarly shaped competing distributions helps us ensure that our results are valid.

We use the log-likelihood ratio test *ℛ* [29, 51] to compare the power-law distribution to other candidate distributions,

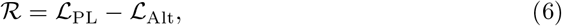

where *ℒ*_PL_ and *ℒ*_Alt_ are the log-likelihoods of the best fits for the power-law and alternative distributions, respectively. This can be rewritten as a summation over individual observations,

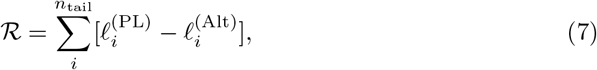

with the log-likelihood of single observed degree values under the power-law distribution, 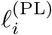, and alternative distribution, 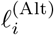, are summed over the number of model observations, *n*_tail_.

If *ℛ >* 0, the power-law distribution is more likely; if *ℛ <* 0, the competing candidate distribution is more likely; if *ℛ* = 0, they are equally likely. Just like with the goodness of fit test, we need to make sure our result is statistically significant (*p <* 0.01). The methodology described here summarizes the methodology introduced by Clauset et al., 2009, and described again in Broido and Clauset, 2018 [29, 30] and more details such as the exact formulas for alternative distributions, and derivation of the *p*-value for *ℛ* can be obtained therein.

### Classifying network scaling

We classify each genomic/metagenomic dataset, as represented by the set of eight network projection types, as having some categorical degree of “scale-freeness” from “super-weak” to “strongest”. This classification scheme was introduced by Broido and Clauset, 2018 [30] in order to compare many networks with different degrees of complexity, and the definitions below were extracted from therein:

- Super-Weak: For at least 50% of graphs, none of the alternative distributions are favored over the power law.

The four remaining definitions are nested, and represent increasing levels of direct evidence that the degree structure of the network data set is scale free:

- Weakest: For at least 50% of graphs, the power-law hypothesis cannot be rejected (*p ≥* 0.1).
- Weak: The requirements of the Weakest set, and there are at least 50 nodes in the distribution’s tail (*n*_tail_ *>* 50).
- Strong: The requirements of the Weak and Super-Weak sets, and that 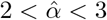.
- Strongest: The requirements of the Strong set for at least 90% of graphs, rather than 50%, and for at least 95% of graphs none of the alternative distributions are favored over the power-law.

Categorizing a network as “Super-Weak” is in effect saying that that network’s degree distribution data is *better* modeled by a power-law fit than alternative distributions. This is independent of whether or not the power-law model is a *good* fit to the data, which is what is what the “Weakest” and “Weak” definitions emphasize. A network may be classified as “Super-Weak” without meeting any of the nested definition’s criteria. Similarly, a network may be classified as “Weak” without meeting the criteria in the “Super-Weak” definition. We believe this framework is a proper way to classify the degree-distributions of biochemical networks, given that there are many different accepted ways to represent biochemical reactions as networks, and each has their pros and cons [32–34].

#### Standard error and correlation

The black error bars on each plot represent 2 standard deviation (2SD) around the sample proportion 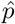 (the height of the bar, which we also refer to as the mean). This is equivalent to 2 standard error around the mean (2SEM), or a 95% confidence interval for the true population proportion *p* (true population mean). Standard deviation was calculated by treating each category as a binomial distribution, meaning the standard deviation is given by:

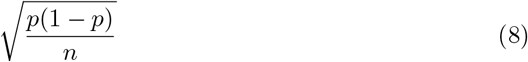

Although the errors for each plot’s categories are calculated independently, there is co-variance between many of them. This is especially true for the right column of **Fig. 3**, where all bars of a color total to a fixed number of datasets, with each dataset falling into one of the 8 network projection type bins. Because of this, we also calculated the correlations between each network projection type, across both individuals and ecosystems (**Fig. S7**). The correlation matrices were calculated by using the pandas function `DataFrame.correlation(method=‘pearson’)` on a matrix of binomially distributed True/False values representing whether each dataset passed or failed specific scale-free criteria for *p*-value, tail size, or power-law exponent value (*α*), for each network-projection.

### Classifying levels of biology using degree distribution data

We used two different statistical methods, multinomial regression and random forest classifiers, in conjunction with the scale-free classification scheme above in order to test if individuals and ecosystems were distinguishable based on their degree distribution characteristics.

#### Multinomial regression

For our multinomial regression, the response class is the biological level (individual or ecosytem), and a single network or statistical measure is the dependent variable. In order to control for over fitting the training data was composed of an equal number of samples from each level. The number of networks used for training data was chosen to be equal in size to 80% of all ecosystem projections, because there were less ecosystem datasets used than individual datasets. This corresponded to 80% of 6280 networks (of all projection types), or 5024 networks. The model was tested on the 20% of the data that it was not trained on. This process was repeated 100 times and the average model error is reported in the results and **Fig. 5**, left columns. The `multinom` and `predict` functions from the R-package `nnet` were used to do the multinomial regression.

#### Random forest classifiers

We used a random forest to attempt to classify networks as falling into the category of individuals or ecosystems. In the first scenario, we used 11 predictors: power-law alpha value (*α*); log-likelihood result from power-law vs. exponential (*dexp*); log-likelihood result from power-law vs. log-normal (*dln*); log-likelihood result from power-law vs. power-law with exponential cutoff (*dplwc*); log-likelihood result from power-law vs. stretched exponential (*dstrexp*); the network mean degree (*< k >*); network node size (*n*); degree distribution tail size (*n*_*tail*_); network edge size (*n*_*edges*_); the *p*-value of the goodness-of-fit test for the power-law model (*p*); and cutoff degree value for network tail (*x*_*min*_). In the second scenario, we repeated the random forest without the three predictors which can be directly used to quantify the size of a network (*n, n*_*tail*_, and *n*_*edges*_). In the third scenario, we repeated the random forest without the three predictors on each network projection type independently. For each scenario, we randomly split our data in two halves: one for training, and one for testing (for the third scenario, each training and testing set is 1/8 as large as for the first two scenarios, since we run the classifier on each network projection type independently). In all scenarios, we use the `randomForest` function from the R-package `randomForest` for classification. Three features were used to construct each tree (`mtry=3`), which is 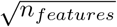, with 100 trees generated each time (enough time for the out-of-bag, or OOB, estimate of the error rate to level off).

## Acknowledgments

We thank the Emergence@ASU team (especially Doug Moore, Cole Mathis, and Jake Hanson) for feedback through various stages of this work.

## Supporting Information

### S1 Fig

#### How alternative distributions compare to the powerlaw across each network projection type

The proportion of network projections, across all datasets, that favor either the power-law distribution (1.0), an alternative distribution (−1.0), or are inconclusive (0.0). Each row shows a different network projection type. Each column is a different distribution with which the power-law is being compared to. From left to right is the exponential; log-normal; stretched exponential; and power-law with cutoff. Dashed line is constant at proportion = 0.5 across all subplots. Red bars indicate individual-level networks, and blue bars indicate ecosystem-level networks.

### S2 Fig

#### Correlations between network projections which meet scale-free criteria

Correlation matrix heatmaps show type how different types of network projections correlate in their proportions of networks which meet some scale-free criteria. Rows are for each of the different scale-free criteria (*p*-value, *n*_*tail*_ and *α*), and columns are for individual and ecosystem-level networks. Heatmaps show the correlation between values for each projection type, where the values are of the proportion of networks which meet the scale-free threshold criteria of: *p* ≥ 0.1 (top row); *n*_*tail*_ *>* 50 (center row); 2 *< α <* 3 (bottom row). Values from projections of a network’s LCC and entire graph are highly correlated. All types of unipartite compound networks tend to be correlated. Values across many other projection types are barely correlated for the *p*-value and *n*_*tail*_ criteria. Ecosystems tend to show more correlation, across all projection types, than individuals.

### S3 Fig

#### Predicting individuals and ecosystems from degree distribution data using multinomial regression

Each subplot shows the accuracy of using a particular network or statistical measure to predict whether that network data came from an biological individual or ecosystem. The subplots in the right column are the accuracy of using a measure after being normalized to network size. Unsurprisingly, network size is by far the best way to accurately predict whether data comes from an individual or ecosystem (left blue star). Once normalized to size, whether or not a degree distribution favors an exponential fit compared to a power-law fit becomes a decent predictor (right blue star). Subplots measures are: power-law alpha value; log-likelihood result from power-law vs. exponential; log-likelihood result from power-law vs. log-normal; log-likelihood result from power-law vs. power-law with exponential cutoff;

## S1 Table

### Random forest accuracy by network projection type

The predictors used in the random forest are the same predictors used in the multinomial regression: power-law alpha value; log-likelihood result from power-law vs. exponential; log-likelihood result from power-law vs. log-normal; log-likelihood result from power-law vs. power-law with exponential cutoff; log-likelihood result from power-law vs. stretched exponential; the network mean degree; network node size; degree distribution tail size; network edge size; the *p*-value of the goodness-of-fit test for the power-law model; cutoff degree value for network tail. See methods for description of network projection types.

The predictors used in the random forest are the same predictors used in the multinomial regression: power-law alpha value; log-likelihood result from power-law vs. exponential; log-likelihood result from power-law vs. log-normal; log-likelihood result from power-law vs. power-law with exponential cutoff; log-likelihood result from power-law vs. stretched exponential; the network mean degree; network node size; degree distribution tail size; network edge size; the *p*-value of the goodness-of-fit test for the power-law model; cutoff degree value for network tail. See methods for description of network projection types.

## Notes

### Competing Interest Statement

The authors have declared no competing interest.

